# Phage-mediated just-in-time circuit amplification and delivery for recombinant expression of toxic proteins

**DOI:** 10.1101/2025.08.31.673379

**Authors:** Leonard E. Bäcker, Flo Baert, Kevin Broux, Dries Oome, Kenneth Simons, Steffen Waldherr, Kristel Bernaerts, Abram Aertsen

**Affiliations:** Department of Microbial and Molecular Systems, Faculty of Bioscience Engineering, KU Leuven, Leuven, Belgium; Department of Chemical Engineering- (Bio)chemical Reactor Engineering and Safety, Faculty of Engineering, KU Leuven, Leuven, Belgium; Current address: Department of Functional and Evolutionary Ecology, Faculty of Life Sciences, University of Vienna, Vienna, Austria

## Abstract

In order to address outstanding challenges in bacterial bioreactor bioprocessing, we have developed an approach that alleviates the need for chemical inducers and selective agents (such as antibiotics), and mitigates leaky expression of the production circuit. More specifically, temperate bacteriophage Lambda (λ) was engineered as a chassis to just-in-time amplify and deliver a production circuit to a wild-type *Escherichia coli* population. Since the expression of the production pathway is engineered to be dependent on λ’s central regulatory switch towards the lysogenic state, λ will first lytically amplify in the background without significantly altering overall bioreactor growth dynamics or gene expression. Only when λ eventually starts outnumbering the host population, its natural switch to lysogeny will actively install and trigger the production circuit. As such, the λ chassis serves as an intrinsic self-amplifying inducer that meanwhile rules out any leaky expression. The impact of the initial phage-to-host ratio on population dynamics and production efficiency was assessed experimentally and through ODE modeling, revealing that it can be used to fine-tune the trade-off between gene dosage and biomass conversion into virions. We demonstrate that this leakproof approach is particularly promising for the expression of toxic proteins such as Benzonase, a highly potent DNase and RNase.

## Introduction

Recent advancements in bioprocessing have brought forward various applications of microbial cell factories for the production of a wide variety of biochemical products. These products range from simple metabolites (e.g. alcohols or organic acids) that can provide important substitutes for petro-chemically derived commodities (1), to complex functional molecules that could not be synthesized otherwise (e.g. therapeutic compounds, enzymes or structurally challenging hydrocarbon derivatives) (2-4). However, while bioprocessing is an increasingly important cornerstone for sustainable development and health (5-7), several challenges still need to be addressed to improve its efficiency and economic viability (8,9).

Indeed, aside from engineering and optimizing the gene or pathway of interest itself, its proper accommodation and timely expression within the cell is important as well (10). In this context, optimal bioreactor yield and productivity are typically achieved by clearly separating microbial growth and production phases, which essentially requires that expression of the engineered gene, or pathway, is fully repressed during the growth phase, and only activated after reaching a sufficiently high biomass density (11). In current applications, the switch from the prior growth to the subsequent production phase is typically brought about by the addition of a chemical inducer (12-14). However, economic cost analyses for the production of recombinant proteins in bioreactors have identified inducers (such as the most commonly used IPTG) as one of the largest raw material cost factors (14,15). While cheaper inducer sources have been explored (16-18), engineered autoinduction mechanisms are meanwhile emerging as an alternative. These latter mechanisms depend on autonomous sensing of cell density (via quorum sensing) or of carbon or phosphate depletion as a cue to transit to the production phase (19,20).

Aside from the actual production trigger, another important issue stems from the burdensome leaky expression of the gene(s) of interest already during the growth phase, with the burden being aggravated when the expressed cargo is toxic for the cell (21-23). Indeed, this tends to select for the enrichment of so-called escape mutants that manage to consume substrate without contributing to production of the desired compound (24). The competitive growth advantage of these non-producing subpopulations of escape mutants becomes particularly detrimental during bioreactor scale-up as more generations of growth are required before target biomass densities for production are achieved (25). While some attempts have been made to stabilize production pathways via the use of product addiction mechanisms, their use is limited to the availability of biosensors for the respective product and/or has inherent design constrains that need to be balanced to accommodate the addiction without fitness burden (26,27). Another popular strategy for improved pathway stability is to move production genes from plasmids to the chromosome. However, while lowering the copy number would indeed help in reducing leaky expression, for yields to be comparable to plasmid-based productivity, generally multiple chromosomal copies of the pathway are required (28,29).

To address both these inducer cost and leaky expression issues, we decided to explore the lytic-lysogenic decision making of a temperate phage as a means to trigger and drive the production phase. More specifically, we engineered temperate phage Lambda (λ) to deliver the circuit for producing the toxic Benzonase protein to an *Escherichia coli* bioreactor population just-in-time for the production phase. The resulting strategy is leakproof and does not rely on the addition of antibiotics or costly inducers.

## Materials and Methods

### Strains, media, and cultivation

For all experiments in this study, *Escherichia coli* K12 MG1655 (further referred to as MG1655), Ur-λ, and derivatives thereof have been used (see Table S1). For cloning of Gibson assembled plasmids *E. coli* DH5α was used as an intermediate host. The strain Ur-λ, was kindly provided by Jay Hinton (University of Liverpool, UK). Lysogeny Broth (LB; 10 g/L tryptone, 5 g/L yeast extract, 5 g/L NaCl; with 1.5% agar added for agar plates) was used for standard cultivation, while LBMM (LB supplemented with 0.2% maltose and 10 mM MgSO_4_) was used whenever the culture was to be used for λ infections. Where needed for genetic engineering purposes, the medium was supplemented with the following antibiotics at the indicated final concentrations: ampicillin (100 µg/mL), chloramphenicol (30 µg/mL), kanamycin (50 µg/mL), tetracycline (20 µg/mL).

All strains were aerobically grown on an orbital shaker at 37°C, with the exception for strains harboring the pKD46 (30) or pCP20 (encoding FLP recombinase (31)) plasmid, which were grown aerobically at 30°C. For stationary-phase cultures, cells were grown for 16-18 h, while for exponential-phase cultures, stationary phase cells were diluted 1/1,000 in fresh medium and grown for an additional 3 h.

### Phage preparations

For the preparation of λ phage stocks, plate lysis was used. First soft-agar (SA) plates were prepared by mixing ca. 10-15 mL SA (10 g/L tryptone, 5 g/L yeast extract, 5 g/L NaCl, 0.5% agar supplemented with 10 mM MgCl_2_ and 1 M CaCl_2_) with 100 µL of a MG1655 overnight culture. Subsequently, the supernatant of an MG1655 λ lysogen was streaked with a loop on the solidified SA plates. After overnight incubation, plaques were excised with a syringe and resuspended in 1 mL suspension medium (SM; 5.8 g/L NaCl, 2 g/L MgSO_4_ × 7H_2_O and 50 mM Tris Cl adjusted to pH 7.5) followed by sterile filtration. Next, tubes were prepared by adding 8 mL molten top agar (TA, 10 g/L tryptone, 2.5 g/L NaCl, 5 g/L agar supplemented with 10 mM MgCl_2_ and 1 mM CaCl_2_) to ca. 0.5-1 mL of exponentially growing MG1655, and 100 µL of serial diluted phage suspension. After brief mixing the agar suspension was distributed on bottom agar (BA, 10 g/L tryptone, 2.5 g/L NaCl, 10 g/L agar) plates preheated to 37°C. The solidified plates were incubated at 37°C overnight.

Plates showing confluent lysis were harvested by first pipetting 2 mL of SM onto each plate followed by chopping up of the TA and collecting the SM and TA mix in a 50 mL Falcon tube. Subsequently, 100 µL of chloroform was added to the tube, followed by rigorous vortexing. The agar slurry was then centrifuged for 10 min at maximum speed, after which the supernatant was gently decanted into a fresh tube to yield the phage stock.

Phage suspensions were titered by mixing 100 µL of an MG1655 overnight culture with 100 µL serially diluted phage suspension and ca 10-15 mL SA on a Petri dish, and determining the plaque count after overnight incubation at 37°C.

### Strain construction

Engineering of the Ur-λ chassis was performed in an MG1655 *recA*^*-*^ Ur-λ lysogen, in which the host *recA* was replaced with a *frt-kmR-frt* cassette to allow *tetA-sacB* counterselection protocols (32) while avoiding prophage curing. Genetic constructs were created via Gibson assembly (33), and engineered within the λ prophage via Lambda Red recombineering (30). The Ur-λ *lom* gene was first replaced by a P1 *parS* sequence, and this strain was considered as the parental stain for any further engineering (referred to as λ from hereon). Subsequently, the λ *rexAB* genes were replaced with the gene encoding T7 RNA Polymerase (yielding λ_*T7*_). From hereon, λ_*T7*_ was equipped with a plasmid (pBR-Lys) bearing the T7 Lysozyme transcribing region from pLysE (34) prior to any further genetic engineering to quench T7 RNA polymerase activity. Subsequently, the λ_*T7*_ *b2 region* was replaced (while leaving in *attP*) with the gene encoding an interrupted PelB-tagged Benzonase (yielding pre-Benzo-λ). The sequence of the PelB-tagged Benzonase was a kind gift from Prof. Rob Lavigne (KU Leuven, Belgium), with the PelB-tag promoting the translocation of Benzonase to the periplasm (35) where it folds into it active form (Lammens et al. in preparation). The gene encoding the PelB-tagged Benzonase was disrupted by a 2.7 kb cassette taken from Näsvall, 2017 (36), comprising a chloramphenicol resistance gene (*cmR*) and a *I-SceI* restriction site, flanked with inverted repeats and short direct repeats.

Since a functional *recA* gene is required to allow for the spontaneous induction of lysogens, we cloned the native MG1655 *recA* gene onto a pACYC184 vector yielding pACYC-RecA. This plasmid was transformed into the engineered MG1655 *recA*^*-*^ pre-Benzo-λ lysogen to yield plaques for the production of phage stocks of the resulting Benzo-λ chassis.

### Phage infection experiments

For infections under batch cultured conditions, exponentially growing MG1655 populations were normalized to an OD_600_ of 0.004 in LBMM (corresponding to ca. 1.6×10^6^ CFU/mL), and either 50 mL were filled into 250 mL Erlenmeyer flasks or 200 µL were filled into flat bottom 96-well plates (Greiner Bio One). The populations were then spiked with phage stock to obtain defined MOIs, followed by incubation at 37°C in an orbital shaker (for flasks) or a plate reader (Multiskan FC, Thermo Scientific; for well plates).

### Microscopy

Fluorescence microscopy images were obtained by placing cells, stained with 1 µL/mL DAPI, between agarose pads (1.5% agarose) and a cover glass using Gene Frames (Life Technologies) and imaged using a Ti-Eclipse inverted microscope (Nikon, Champigny-sur-Marne, France), equipped with a 60× Plan Apo λ oil objective, a TI-CT-E motorized condenser and a Nikon DS-Qi2 camera. DAPI was imaged with a triple dichroic (475/540/600 nm) emission filter, and a SpectraX LED illuminator (Lumencor, Beaverton, USA) as a light source.

### Benzonase quantification

For the quantification of Benzonase production, MG1655 populations were centrifuged for 10 min at 20,000 × *g*, after which the cells were resuspended in a lysis buffer (10 mM Tris-HCl pH 7.2 and 150 mM NaCl) and constantly kept on ice. Cells were lysed using an immersion sonicator (VCX30, Sonics & Materials, Newtown, CT, USA) for a total of 2 min (pulses of 20 s at 30% amplitude with 20 s pauses in between). The lysate was centrifuged for 10 min at 20,000 × *g*, and the supernatant was recovered. Protein extraction efficiencies were determined for later yield normalization over total extracted cellular protein (Qubit 2.0, Thermo Fisher Scientific). Next, the samples were directly used for Benzonase activity quantification using the DNase alert kit (IDT, Leuven, Belgium) according to the manufacturer protocol. The fluorescence output produced by the kit was quantified via a multiplate reader (CLARIOstar plus, BMG Labtech) and Benzonase titers were calculated according to a calibration curve using Benzonase stocks (Sigma -Aldrich, Saint Louis, MO, USA).

### Infection dynamics model

An ordinary differential equations (ODE) model for phage infection dynamics was developed using the programming language Julia v1.10 (37). The model simulates a well-mixed batch reactor, containing both bacterial cells growing towards an experimentally defined carrying capacity and phages which can infect the bacteria. The bacterial cells are subdivided into compartments based on their current state, such as: susceptible cells, deciding cells, lytic cells and lysogenic cells. The initial inoculum purely contains susceptible cells, which upon infection enter the deciding compartment, and remain there for a fixed decision period. Subsequently, they enter either the lytic compartment or the lysogenic compartment, depending on the perceived MOI during their stay in the initial deciding compartment. Cells that enter the lytic compartment will burst after a variable duration called the lytic delay, mimicking biological variance in latent periods. While lysogenic cells will stably remain in the lysogenic compartment and grow at an equal rate as the susceptible cells. The corresponding model equations and further documentation can be found in the supplementary information.

## Results

### Design and construction of a phage chassis for just-in-time gene delivery

We envisaged a strategy whereby a temperate phage is first engineered to carry the cargo gene(s) of interest under control of a lysogeny-specific promoter, and is administered under low phage-to-host ratios to a growing bioreactor population. The lytic amplification of this phage will consume a small fraction of this host population to eventually produce a high phage-to-host ratio that subsequently lysogenizes the remainder of the bioreactor population, thereby “just-in-time” providing the host cells with the cargo gene(s) for the production phase. As such, the engineered phage serves as an intrinsic self-amplifying inducer and meanwhile rules out any leaky expression since cells do not yet harbor the cargo gene(s) of interest during their growth phase (Fig. 1).

**Figure 1.**
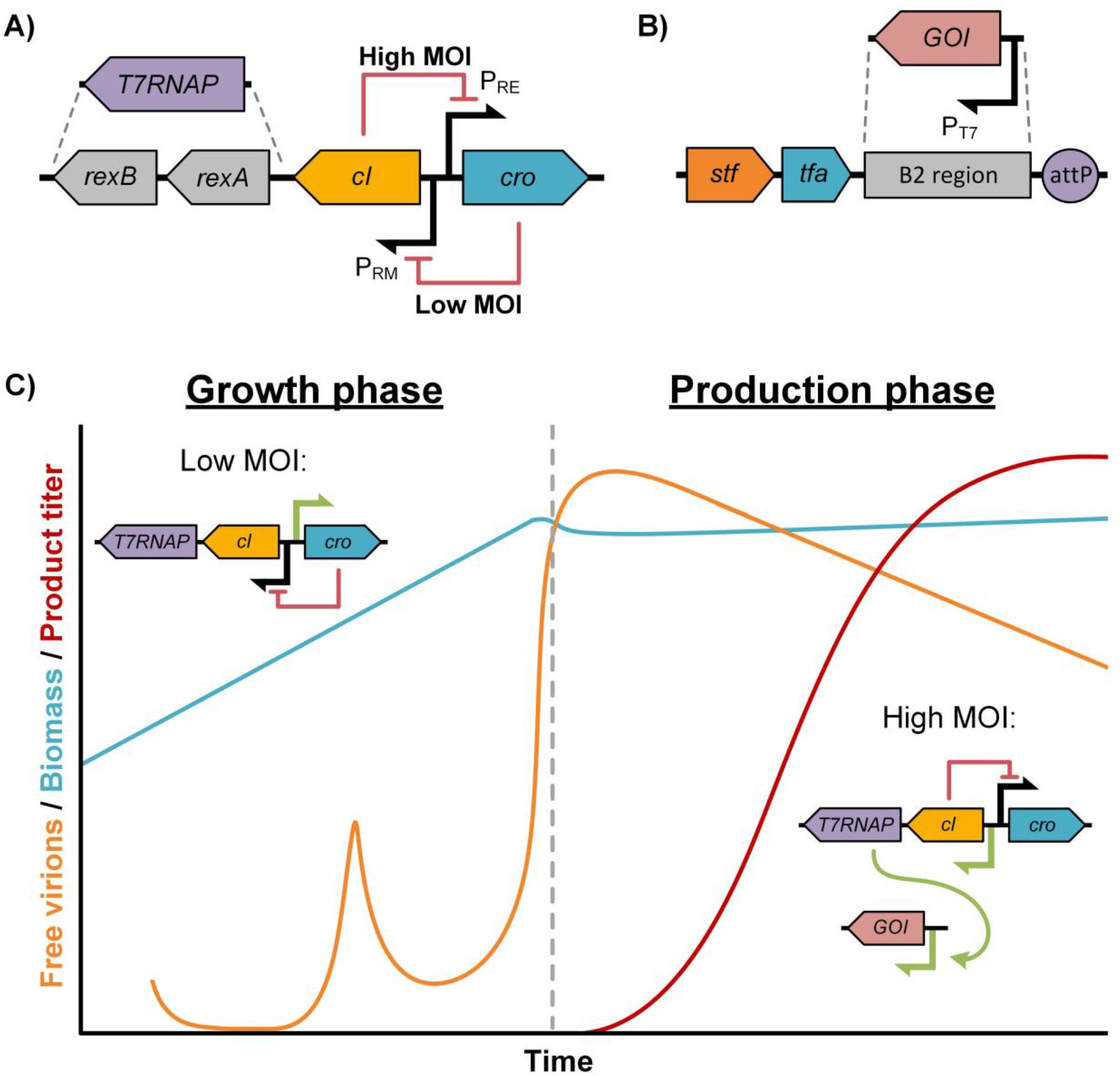
Overview of the genetic modifications in the engineered phage λ chassis. (A) The *rexAB* genes located directly downstream of *cI* have been replaced by the gene encoding T7 RNA polymerase (*T7RNAP*), therefore coupling its expression to λ’s transition from lytic to lysogenic propagation. (B) Genes located in the *b2* region on the λ genome are being replaced by respective genes of interest (GOI) under control of a T7RNAP responsive P_T7_. (C) Illustration of the overall population and circuit dynamics related to infection with the engineered λ chassis, with initial lytic amplification during the bioreactor growth phase eventually ushering the production phase upon reaching a high MOI.

For this study, we opted for *Escherichia coli* as a host that is widely used in bioprocessing, and for temperate phage Lambda (λ), of which the genetics are well understood (38-40). As such, we coupled the shift towards cargo production to the λ P_RM_/P_R_ toggle switch that is central to its native lysis/lysogeny decision making (41). More specifically, we transcriptionally fused the gene encoding T7 RNA polymerase (T7RNAP) to *cI* expression by replacing the non-essential *rexAB* genes (40,42), yielding the intermediate construction strain referred to as λ_*T7*_ (Fig. 1A). During the initial lytic propagation at a low multiplicity-of-infection (MOI), *cro* is transcribed from P_RM_’s opposing promoter P_R_, thereby blocking *cI* and thus *T7RNAP* expression. However, when λ_*T7*_ eventually outnumbers its host (reaching a high MOI), CII will naturally be produced at sufficiently high levels to temporally inhibit lytic functions and cause a spike of *cI* and *T7RNAP* expression (40,43,44). This transcriptional burst is followed by a switch to the CI-silenced lysogeny state (40) in which CI (and therefore also T7RNAP) expression is further maintained. As a result, an expression pathway driven by a T7RNAP responsive promoter (P_T7_) will directly be coupled to a switch towards lysogeny and initiate the production stage (Fig. 1C).

In order to minimally alter the native phage biology, this expression pathway should be transcriptionally isolated and ideally be variable in size without altering the phage phenotype. Therefore, we have identified the non-essential *b2* region (leaving in *attP*) (45,46) between the major early (pL driven) and late (pR’ driven) operon as a landing site for the P_T7_ expressed gene(s) of interest (GOI) flanked by very strong terminators (47,48) (Fig. 1B). Nevertheless, a leftward orientation is recommended to avoid transcriptional interference from the pR’ operon during lytic cycles.

### Lambda-mediated delivery of a highly toxic cargo

An important benefit of avoiding leaky expression is that highly toxic genes, which are inherently unstable in their production host, do not pose any fitness burden or selection pressure. As a proof-of-concept, we therefore decided to express PelB-tagged Benzonase, which readily translocases to the periplasm where it accumulates in its active form. Expression of this powerful DNase (and RNase) can nevertheless be toxic to the host chromosome due to residual activity of cytoplasmically produced Benzonase and/or the leakage of periplasmically folded Benzonase back into the cytoplasm (49,50). In fact, the toxicity of the PelB-tagged Benzonase (further referred to as Benzonase) already became clear by the inability to fit it downstream of the P_T7_ promoter in the MG1655 *recA*^*-*^ λ_T7_ lysogen construction strain. Since *recA* deficient strains are hypersensitive to DNA damage (51), this further suggests the presence of benzonase activity in the cytoplasm.

To avoid any leaky toxicity in the construction strain, we devised a cloning strategy based on the recently developed approach of using long inverted repeats (IRs) of DNA in combination with short direct repeats (DRs) as a high frequency recombinational hotspot (36). More specifically, we equipped an MG1655 *recA*^*-*^ λ_*T7*_ lysogen with a P_T7_-driven *benzonase* construct disrupted by a 2.7 kb cassette comprising a chloramphenicol resistance gene (*cmR*) and a *I-SceI* restriction site, which are flanked with IRs (encoding *amilCP*) and DRs (Fig. 2A). Since the hairpin structure formed by the IRs brings the DRs in close proximity to each other, the latter are prone to recombine and functionally reconstitute the Benzonase encoding gene during prophage induction (Fig. 2A). An MG1655 host expressing the *I-SceI* restriction endonuclease was then used as a plaquing host to select for λ_*T7*_ derivatives that successfully lost the interrupting cassette. However, this counter-selection host appeared not to be essential, likely because the length of the cassette already attenuated proper packaging, automatically favoring recombined phage progeny. Via this procedure, λ_*T7*_-P_T7_-*benzonase* plaques could be readily isolated and purified carrying a reconstituted *benzonase*, leading to the clone further referred to as Benzo-λ.

**Figure 2.**
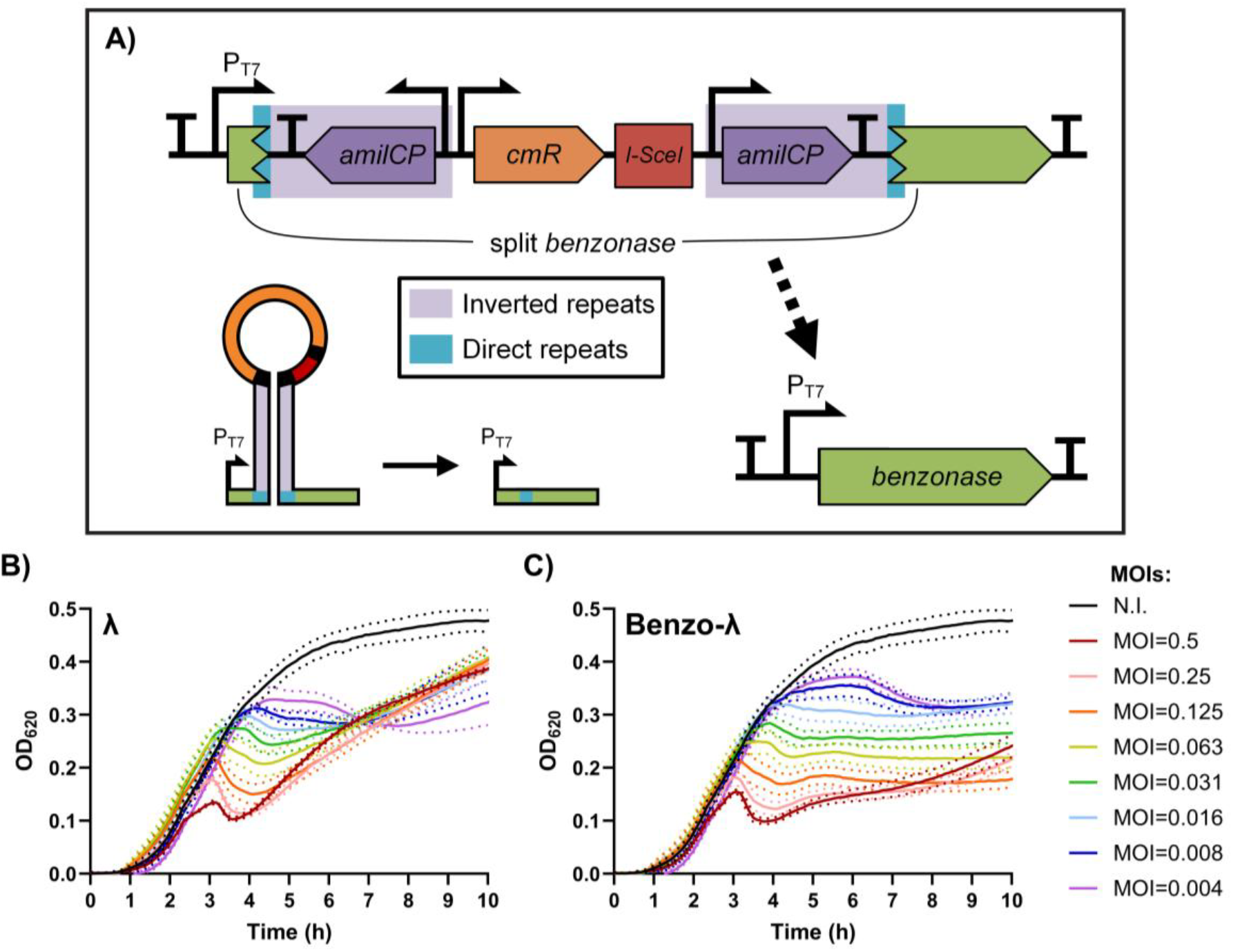
Cloning strategy and initial characterization of Benzo-λ. (A) A *benzonase* ORF (shown in green) under control of a P_T7_ promoter is disrupted in vitro with a cassette for positive and negative selection. The 30bp direct repeats (DRs) of the *benzonase* ORF are highlighted in teal, while operons for the expression of the purple chromoprotein *amilCP* (shown in dark purple) making up the inverted repeats (IRs) are highlighted in purple. Flanked by the repeat regions are a chloramphenicol resistance gene (*cmR*, shown in orange) and an *I-SceI* restriction site (shown in red). The inverted repeats form a hairpin that facilitates recombination into the functional *benzonase* ORF. (B, C) Comparisons of *E. coli* MG1655 cultures infected at different MOIs (0.5-0.004) with either parental λ (B) or engineered Benzo-λ (C). Means and standard deviations (shown as dotted lines) represent the results obtained from two independent repetitions, both performed in triplicates for each respective condition. N.I. indicates non infected.

### Characterizing Benzo-λ infection dynamics

Since significant parts of the phage genome were deleted and unintended expression of Benzonase during lytic development would be detrimental to phage propagation, we compared the overall infection dynamics of Benzo-λ with its λ parent on wild-type MG1655 over a range of different initial MOIs (Fig. 2B and C). This revealed that the prior lytic infection dynamics (as judged by onset and magnitude of the OD_620_-drop) remained very similar, suggesting that Benzo-λ is indeed not burdened by its extensive engineering or by potential leaky expression of Benzonase during lytic propagation.

Additionally, in contrast to λ lysogens, Benzo-λ lysogens clearly stopped growing during the lysogenization phase following the OD_620_-drop, indicating the intended production of the toxic Benzonase. Interestingly though, we did not observe any further drop in the OD of Benzo-λ infected cultures, suggesting that the metabolic burden of full Benzonase production (and the DNA damage it causes) hampers proper lytic activation of the Benzo-λ prophage (52,53).

More closely monitoring MG1655 Benzo-λ infection dynamics (with starting MOI of 0.06) via both OD_600_ measurements and viability plating indeed revealed a massive (2-3 log) loss of viability compared to λ infections once high MOI conditions were reached, indicating that the vast majority of the population was properly converted into Benzonase producers (Fig. 3A-C). When samples were subsequently taken for DAPI staining at 2 h, 4 h and 6 h after initial infection (Fig. 3D-F), it could indeed be observed that from 4 h ca. 60% of Benzo-λ lysogens did not appear to have any stainable DNA left (Fig. 3G). The nucleoid of the remaining cells also appeared less intensely stained and diffuse compared to both λ-infected or non-infected controls, suggesting a significant degree of DNA degradation. Please note that, as seen by the very bright staining of λ infected hosts, incoming DNA from infecting phages seems to contribute to the observed DAPI staining intensity. As such, the relatively high MOI conditions at 4 and 6 h are likely to have contributed to the partial DAPI staining in Benzo-λ infected cultures due to recent superinfection events. This further underscores that, although PelB-tagged Benzonase translocates to the periplasm, the high expression level nevertheless increases Benzonase activity to lethal doses within the cytoplasm.

**Figure 3.**
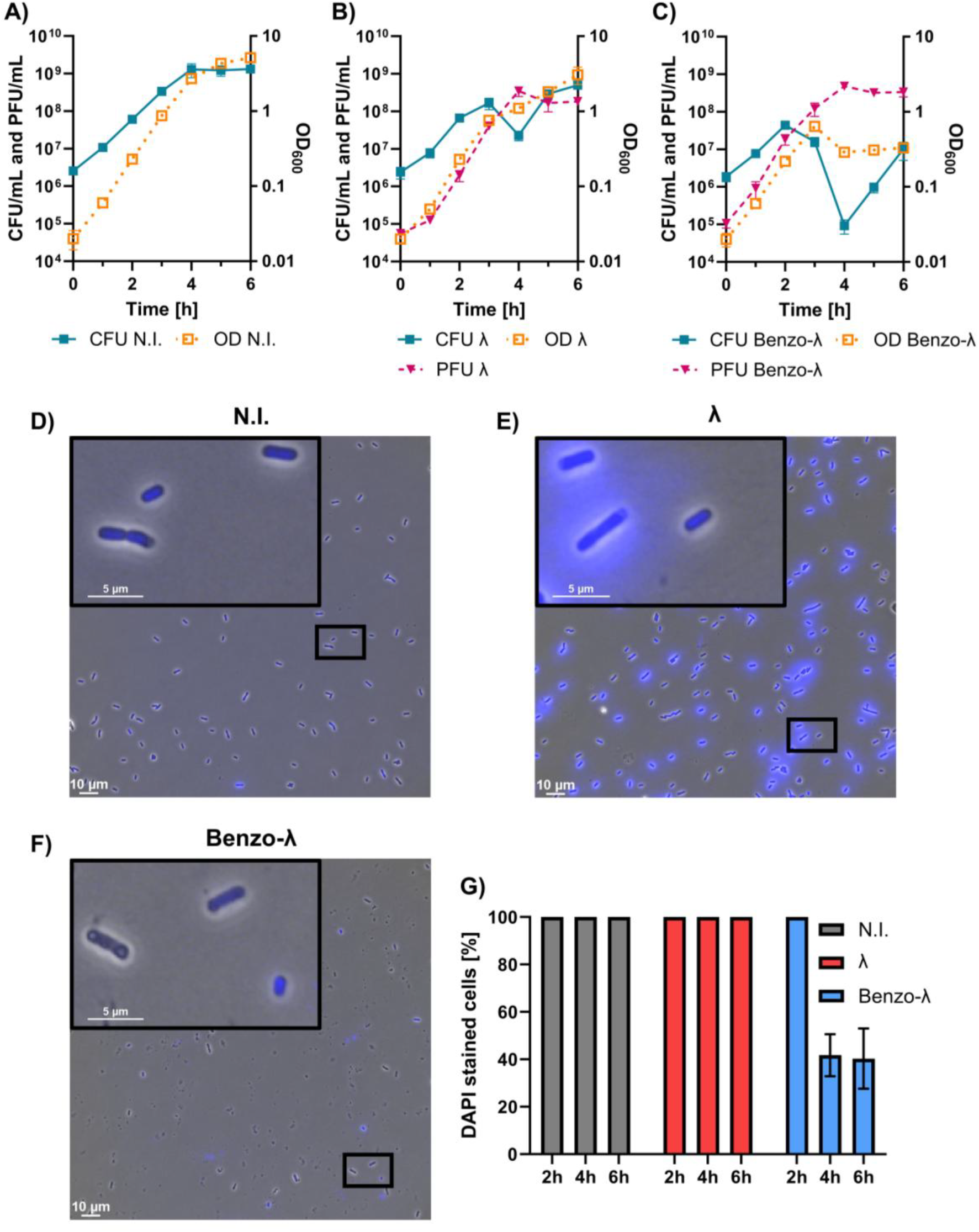
Impact of engineered phage Benzo-λ infections on host cells through the production of Benzonase. *E. coli* MG1655 cultures were (A) non-infected (N.I. controls) or identically treated but infected with a low MOI (0.06) of either λ (B) or Benzo-λ (C) and monitored for viable cell count (lines representing CFU/mL), viable phage count (dashed lines representing PFU/mL), and OD_600_ (dotted lines). For (A) and (B) means and standard deviations represent the results of three independent replicates (n=3), while for (C) three independent phage stocks were assessed in three independent replicates respectively (n=9). Representative fluorescence microscopy images of DAPI stained samples, with (D), (E) and (F) respectively being taken from the cultures shown in panel (A), (B) and (C) after 6 h of cultivation. The fluorescence intensity of the images can be directly compared and was analyzed to calculate the percentage of cells (952 ± 504 per data point) that showed fluorescence significantly above background levels (G) with standard deviations shown in bars.

The two log increase in viable cell count in the Benzo-λ infected cultures from 4-6 h (Fig. 3B) seemed peculiar, suggesting that a small fraction of cells did not yet encounter Benzo-λ due to improper mixing in the shake flasks and/or an overabundance of lysogenic cells titering away the free Benzo-λ particles and thus delaying infection of these naïve cells.

### Tuning Lambda-mediated delivery to maximize Benzonase production

Since the initially added amount of phages can heavily influence the infection dynamics (Fig. 2B and C) and thus the timing of lysogenization and productivity, we followed Benzo-λ infections starting from 18 different initial MOIs (0.41-0.00025; Fig. S1). Interestingly, this revealed that the fraction of cells that are lost due to lytic phage propagation (until high MOI infections result in lysogenization) shows an oscillating pattern in function of the applied MOI (Fig. 4A). To see how these dynamics impact phage-mediated productivity, we quantified the yield of intracellularly produced Benzonase for selected initial MOIs that correspond to the peaks and valleys of the biomass loss trend. Surprisingly, however, experimentally observed Benzonase yields showed a peculiar independence regarding high or low biomass losses (Fig. 4A).

**Figure 4.**
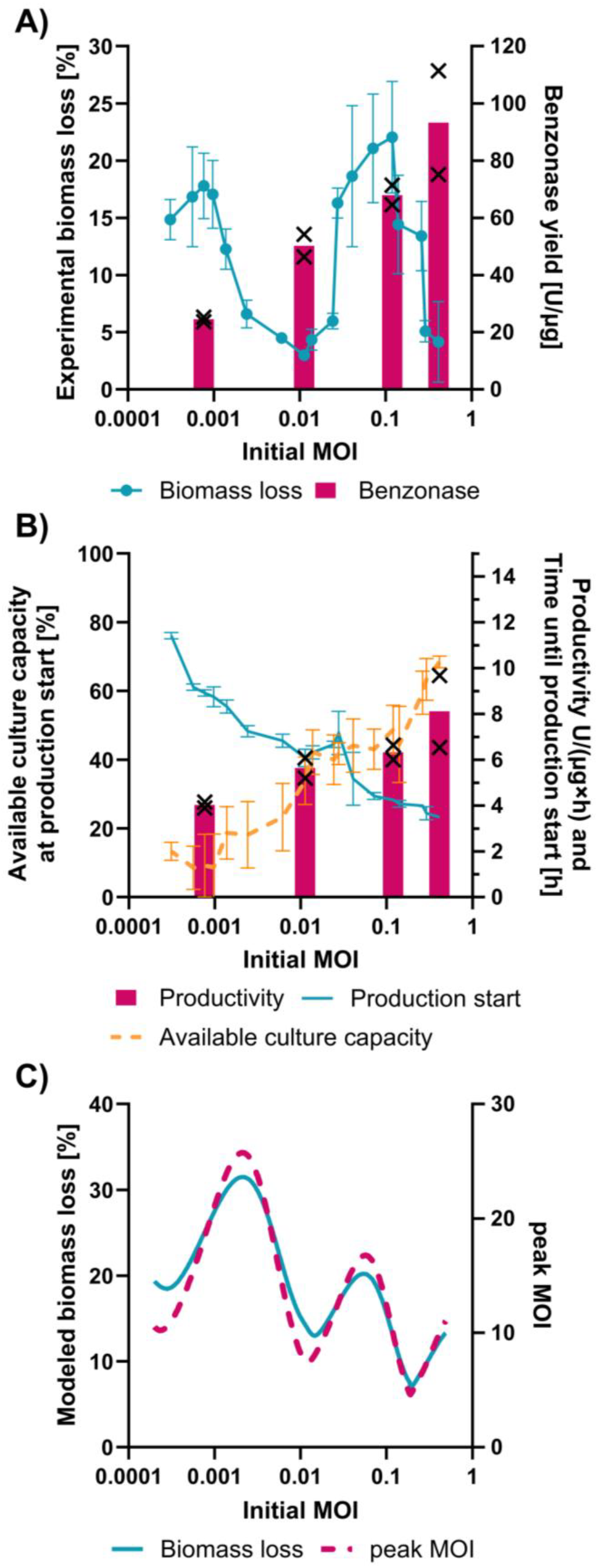
Lysogenic conversion dynamics to initiate Benzonase production in relation to the initial phage input. (A) Relative loss in biomass at lysogenic conversion for Benzo-λ infections (Fig. S1) calculated in relative percentage of the minimum OD_620_ after lysogenization compared to the maximum OD_620_ before lysogenization, as well as the Benzonase yield for the indicated MOIs. (B) Production start for Benzonase expression, determined as the time point the minimum OD_620_ during lysogenization is reached. Based on the production start and the end of incubation after 15 h, the productivity is calculated as an hourly rate. The available culture capacity is given as the percentage fraction of the maximum OD_620_ before lysogenization relative to the total maximum OD_620_ of the non-infected controls. The results shown in (A) and (B) represent the averages of three independent replicates, with the exception of the Benzonase values for which three samples have been pooled followed by quantification in technical duplicates. The resulting individual values are respectively indicated by black “x” symbols. (C) Modeling of the infection dynamics shown in panel (A), with biomass loss relative to the biomass at lysogenic conversion (“Biomass loss”, similar to panel (A)), while the highest achieved MOI throughout the respective infection is given as “peak MOI”.

We subsequently modeled λ infection dynamics using the same starting input parameters as used in the experiments of Figure 4A and B. This confirmed the same oscillatory pattern, and further linked it to the MOI at lysogenic conversion (Fig. 4C). In fact, a very low biomass loss is observed when the last lytic cycle (e.g. before phages start outnumbering their host and start lysogenization) creates a sufficiently large overshoot (e.g. MOI∼10) to force most of the remaining cells into lysogeny. However, as indicated by the model, when such a large overshoot is not obtained and MOIs closer to 1 are reached, a large fraction of cells becomes inclined for lytic infection at the cost of higher biomass loss. Interestingly, however, the latter also creates a higher peak MOI due to the additional conversion of biomass into phage particles (Fig. 4C), and as such results in a higher dose of incoming Benzo-λ chromosomes. In turn, this higher intracellular copy number of the Benzonase gene likely boosts the productivity per cell, thereby offsetting the impact of biomass loss on productivity. Therefore, the initial MOI can be adjusted according to whether biomass yield or copy number is the more desired process characteristic.

Finally, the total Benzonase yield appears to strongly correlate with increasing initially imposed MOIs. Lower initially imposed MOIs result in a more postponed lysogenic conversion of the population (Fig 4B, shown as “production start”), which is expected to trivially result in a lower availability of nutrients during the production phase due to medium depletion. Indeed, when converting the peak OD_620_ just before lysogenic takeover into a percentage fraction of the capacity limit of the culture (i.e. maximum OD_620_ in the non-infected controls; Fig. 4B, shown as “available culture capacity”) we can observe a strong correlation with the observed Benzonase productivity trends (Fig 4B).

## Discussion

The development of new bioprocessing approaches that reduce costs and increase yields are important for further maturation of this field. As such, our phage-mediated just-in-time gene amplification and delivery approach has a number of unique benefits. First of all, due to the self-amplifying character of phages, this approach automatically provides its own inducer. As such, it bypasses the use of chemical inducers that are often one of the largest raw material costs in bioprocessing applications (14,15). The cells that meanwhile become lysed to support phage amplification can be readily cannibalized by *E. coli* and thus serve as nutrients for growth and production (54). At the same time, using a phage chassis also bypasses the use of antibiotics that are often used to select for plasmids containing the production circuit. The use of antibiotics in bioprocessing is becoming increasingly disfavored due to costs, decreased selection efficiency during scale-up (especially β-lactam antibiotics) and its links to polar effects (phenotype/burden), while at the same increasingly stricter regulations are adding further pressure to investigate alternatives (55-59).

Secondly, since temperate phage λ strictly separates its lytic from its lysogenic gene expression, the P_RM_-dependent production circuit essentially remains passive cargo during lytic amplification of the phage. This in turn prevents possible leaky expression during the bioreactor growth phase, which depending on the toxicity can cause the emergence of non-productive escape mutants. Upon upscaling of bioreactor populations, take-over by escape mutants can become particularly problematic (25).

Thirdly, via the imposed initial MOI input, the production-triggering lysogenic conversion event can be tuned to occur at the desired time or population density. Moreover, the initial MOI also serves as input to adjust the copy number of delivered production circuits, thereby allowing to tune production beyond the stepwise changes in plasmid copy number offered by different origins of replication (60). An added benefit of just-in-time receiving multiple copies of the production circuit is its robustness to mutations. Indeed, while circuit mutations in conventional expression systems often result in non-productive escape mutants, they become in the current system compensated by a majority of correct production circuits delivered to the cell (hence preventing the cell from becoming a non-producer).

Also, the fact that wild-type hosts can be used with cognate temperate phages opens up the possibility to reprogram host populations which are already established in a nice (e.g. environmental consortia or gut microbiome), which would otherwise require the introduction of engineered strains in those already occupied niches.

Finally, although the expression pathway for Benzonase production used for this proof-of-concept study is relatively small, larger metabolic pathways can in the future be accommodated on our phage chassis as well. Although the engineered λ chromosome should not exceed the 53 kb maximum genome size that can still be efficiently packaged into the phage capsid, many known non-essential λ regions can still be removed (36,38,40). As such, major parts of the non-essential *nin*-region (61,62) and the non-essential *stf/tfa* genes (63) can be deleted to yield up to 6kb extra cargo space, without severely altering the phage phenotype.

In conclusion, our proposed use of synthetically engineered temperate phages to direct bioreactor populations provides a new approach for the efficient expression of (even toxic) proteins in engineered or even wild-type hosts. Indeed, the just-in-time amplification and delivery of expression circuitry alleviates the use of costly inducers and inconvenient antibiotics, and prevents positive selection on non-productive circuit mutations.

## Acknowledgements

This work was funded by doctoral fellowships (11H0323N to L.E.B., 11Q4T24N to Ke.B.) and project grant (G0C5322N) from the Research Foundation-Flanders (FWO-Vlaanderen). The authors would like to thank Rob Lavigne, Vincent De Maesschalck and Maarten Boon for technical assistance in the quantification of Benzonase activity.

## Author Contributions

L.E.B., Kr.B., S.W. and A.A. conceptualized the project. L.E.B., Kr.B., and A.A. designed experiments. L.E.B. and F.B. performed experiments. L.E.B. performed data analysis. D.O. performed the ODE Modeling. L.E.B. and Ke.B. performed image analysis. L.E.B. and A.A. drafted the manuscript and all authors edited the manuscript.

