## Supplemental Data for "Phage-mediated just-in-time circuit amplification and delivery for recombinant expression of toxic proteins"

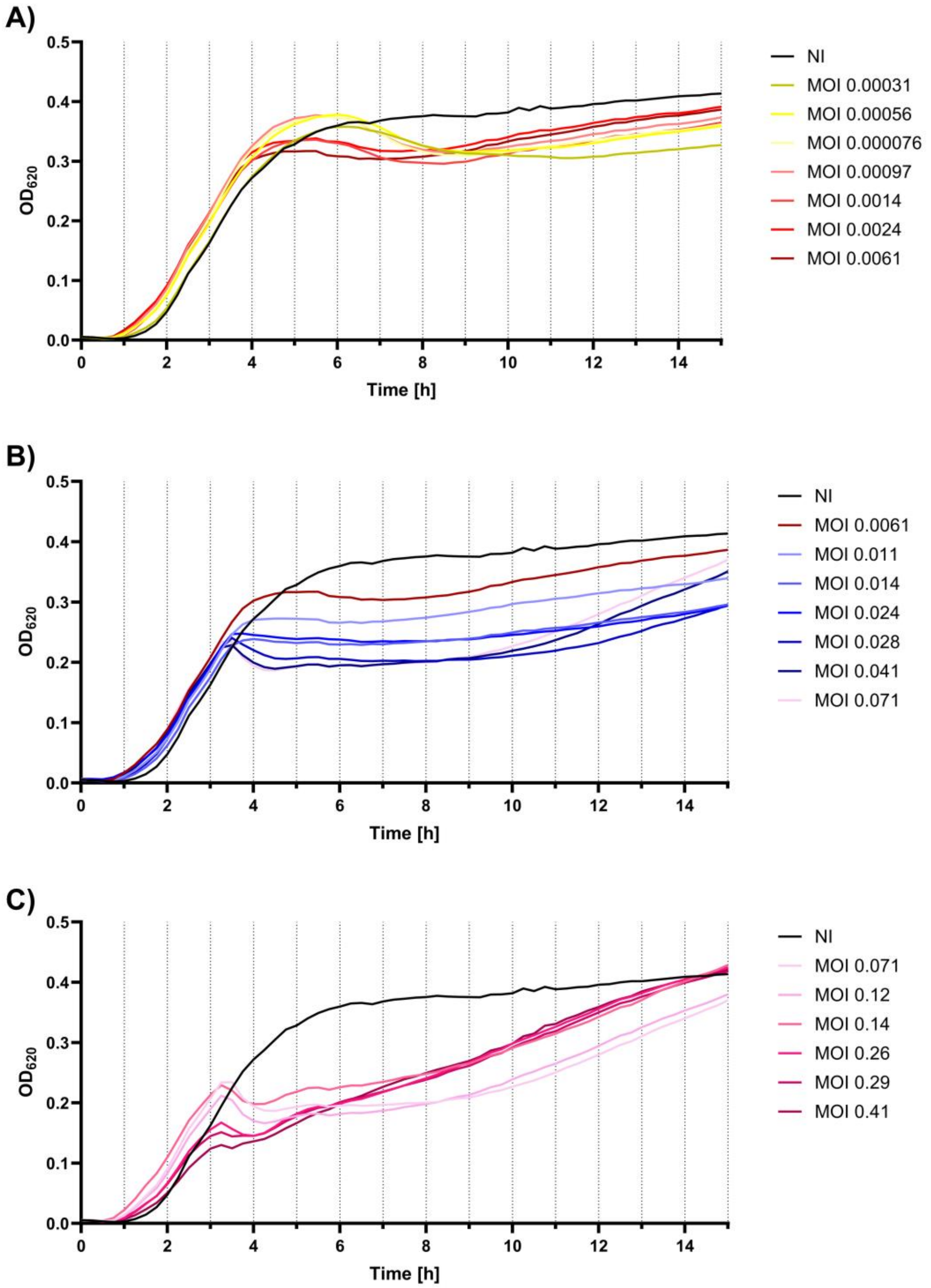


Figure S1: $OD_{620}$ evolution over time of an exponential MG1655 population exposed to 18 different Benzo-λ concentrations (resulting MOIs are indicated). For clarity, each graph is showing 6 different MOIs, as indicated in the legend of (A), (B) and (C). Each line represents the average of three independent replicates. Standard deviations were omitted for the sake of visibility.

Table S1: List of strains used in Chapter 5

| Bacterial strains | Source |
|---|---|
| **MG1655**<br><br>*Escherichia coli* K12 derivative MG1655 wild-type | (1) |
| **DH5α**<br><br>*Escherichia coli* MG1655 derivative with the genotype*:* endA1 hsdR17 supE44 λ– thi-1 recA1 gyrA96 relA1 Δ(lacZYA-argF)U169 φ80lacZ ΔM15 | (2) |
| Intermediate phage construction hosts | Source |
| **MG1655 *recA*$^{-}$**<br><br>*Escherichia coli* MG1655 derivative in which the native recA gene has been replaced with a *frt-kmR-frt* cassette followed by marker curing leaving behind a *frt* scar | This study |
| **MG1655 *recA*$^{-}$ pBR-Lys**<br><br>*Escherichia coli* MG1655 derivative devoid of the native recA gene carrying a plasmid for the constitutive expression of T7-lysozyme | This study |
| **MG1655 *recA*$^{-}$ pBR-Lys pACYC-recA**<br><br>*Escherichia coli* MG1655 derivative devoid of the native recA gene carrying a plasmid for the constitutive expression of T7-lysozyme and a plasmid bearing the native recA locus | This study |
| Bacteriophage strains | Source |
| **Ur-λ**<br><br>Ur-λ wild-type with intact STF genes unlike the common PaPa-λ lab strain. This is the parental strain for all subsequent constructs | (3) |
| **λ**<br><br>Ur-λ derivative with *parS* sequence inserted in place of *lom* | This study |
| **$\lambda_{T7}$**<br><br>λ derivative with a *T7RNAP* inserted in place of *rexAB* | This study |
| **Benzo-λ**<br><br>$\lambda_{T7}$ derivative with a $P_{T7}$-driven *benzonase* construct inserted into the *b2* region | This study |

Table S2: Phage infection model parameters

| Name | Symbol | Value | Units | Description |
|---|---|---|---|---|
| **Carrying capacity** | $K$ | $10^{12}$ | $\frac{Cells}{L}$ | The carrying capacity reflects the total number of bacteria that can be produced based on the (virtual) starting substrate concentration. |
| **Starting biomass concentrations** | $S(0)$ | $1.6 * 10^9$ | $\frac{Cells}{L}$ | The starting biomass is the total number of bacteria in the initial inoculum. All are assumed to belong to the susceptible compartment. |
| **Starting phage to host ratio** | $\frac{P(0)}{B(0)}$ | $[10^{-5}, 10^{-1}]$ | $\frac{phages}{host}$ | The ratio of phages to hosts present in the batch at time $t = 0$. |
| **Maximum growth rate** | $\mu_{max}$ | 2.08 | $\frac{1}{h}$ | The maximum growth rate of the bacteria. |
| **Phage absorption rate** | $\alpha$ | $4 * 10^{-10}$ | $\frac{1}{h}$ | The rate at which phages can absorb to their bacterial hosts. |
| **Burst size** | $b$ | 100 | $\frac{phages}{burst}$ | The number of phages that are released per lysis event. |
| **Decision delay** | $\tau_d$ | 0.5 | $h$ | The time it takes for a bacterium that is infected by at least one phage to decide on the infection outcome. |
| **Mean lytic delay** | $\tau_\mu$ | 0.5 | $h$ | The average time it takes a lytic bacterium to lyse starting from their entry into the lytic compartment. |
| **Variance lytic delay** | $\tau_{\sigma^2}$ | 0.005 | $h$ | The variance of the lytic delay distribution. |

Table S3: Phage infection model variables

| Name | Symbol | Units |
|---|---|---|
| **Susceptible cells** | $S(t)$ | $\frac{Cells}{L}$ |
| **Deciding cells** | $D(t)$ | $\frac{Cells}{L}$ |
| **Lytic cells** | $L(t)$ | $\frac{Cells}{L}$ |
| **Lysogenic cells** | $l(t)$ | $\frac{Cells}{L}$ |
| **Phages** | $P(t)$ | $\frac{Phages}{L}$ |
| **MOI** | $\phi(t)$ | $\frac{Infections}{Cell * \tau_d}$ |
| **Total Produced Biomass** | $B_h(t)$ | $\frac{Cells}{L}$ |

### Equations

The total bacterial biomass is divided into four compartments. The susceptible compartment $S$, the deciding compartment $D$, the lytic compartment $L$ and the lysogenic compartment $l$. The summation of these compartments gives the total current biomass, as given in Equation 1.

$$B(t) = S(t) + D(t) + L(t) + l(t) \tag{1}$$

Equation 2 is used to determine the fraction of lysogenic infections at time $t$. This equation approximates a fit describing the expected fraction of lysogens given the average MOI.

$$\eta(\phi(t)) = 1 - \frac{2^{\phi(t)}}{2^{\phi(t)} + 1} \tag{2}$$

The growth rate is given by Equation 3 and is based on the logistic growth model (4). Growth in the batch slows down as the total produced biomass $B_h$ approaches the carrying capacity $K$.

$$\mu(t) = \mu_{max} \cdot \left(1 - \frac{B_h(t)}{K}\right)$$

( 3 )

The susceptible biomass compartment represents bacteria that are not yet infected by a phage. The bacteria in this compartment can grow/divide depending on the current growth rate calculated by Equation 3. The rate of infection is adapted from Ellis and Delbruck, 1939 (5), and is linearly depended on the phage concentration and bacterial concentration. The rate of change of the susceptible biomass is shown in Equation 4.

$$\frac{dS(t)}{dt} = \mu(t) \cdot S(t) - \alpha \cdot P(t) \cdot S(t)$$

( 4 )

The deciding biomass compartment represents bacteria who have been infected by at least one phage and are thus in the decision phase of infection. Deciding bacteria remain in this compartment until a fixed decision period $\tau_d$ has passed. After which, they are divided into the lytic $L$ or lysogenic compartment $l$ based on the average amount of infections during the decision period $\tau_d$. The rate at which bacteria are leaving this compartment at time $t$ is equal to the rate at which bacteria entered the compartment at time $t - \tau_d$.

$$\frac{dD(t)}{dt} = \alpha \cdot P(t) \cdot S(t) - \alpha \cdot P(t - \tau_d) \cdot S(t - \tau_d)$$

( 5 )

When infected bacteria enter the lytic compartment, they stay there for a variable amount of time $\tau$ until they lyse and release new phages. The rate at which bacteria enter the lytic compartment $\frac{dL_{in}(t)}{dt}$ is given by Equation 6. The lytic delay $\tau$ is assumed to be normally distributed $\sim\mathcal{N}(\tau_\mu, \tau_{\sigma^2})$. By using a delayed differential equation (DDE), the amount of infected bacteria entering the lytic compartment can be calculated for all previous time points $t_h \in [0, t[$. To determine the contribution of each previous time point $t_h$ to the number of lysing bacteria at time $t$, one needs to multiply the value of the probability density function of $\mathcal{N}(t_e + \tau_\mu, \tau_{\sigma^2})$ evaluated at time $t$ with $\frac{dL_{in}(t_e)}{dt}$. The total amount of lysing bacteria at time $t$ is then given by the integral in Equation 7. The rate of change of the lytic biomass is then calculated using Equation 8.

$$\frac{dL_{in}(t)}{dt} = \left(1 - \eta(\phi(t))\right) \cdot \alpha \cdot P(t - \tau_d) \cdot S(t - \tau_d) \tag{6}$$

$$\frac{dL_{out}(t)}{dt} = \int_0^t \frac{dL_{in}(t_e)}{dt} \cdot pdf\left(\mathcal{N}(t_e + \tau_\mu, \tau_{\sigma^2}), t\right) dt_e \tag{7}$$

$$\frac{dL(t)}{dt} = \frac{dL_{in}(t)}{dt} - \frac{dL_{out}(t)}{dt} \tag{8}$$

After the decision period $\tau_d$ bacteria can alternatively enter the lysogenic compartment. The amount of bacteria entering the lysogenic compartment is equal to the amount of bacteria leaving the deciding compartment at time $t$ multiplied by the fraction of bacteria choosing the lysogenic state. Similar to the susceptible biomass, lysogenic biomass can grow/divide depending on the immediate growth rate $\mu(t)$. The rate of change for the lysogenic biomass is given by Equation 9.

$$\frac{dl(t)}{dt} = \mu(t) \cdot l(t) + \eta(\phi(t)) \cdot P(t - \tau_d) \cdot S(t - \tau_d) \tag{9}$$

The rate of change of the phage concentration is depended on the absorption of phages by the biomass and the release of new virions by lysing bacteria, and is given by Equation 10. It is assumed that all biomass, regardless of their current state, can absorb phages. Additionally, each lysis event is assumed to add a fixed amount of phages $b$ to the culture.

$$\frac{dP(t)}{dt} = -\alpha \cdot B(t) \cdot P(t) + b \cdot \frac{dL_{out}(t)}{dt} \tag{10}$$

The MOI at each point in time $t$ is equal to the average amount of infections per bacteria during the period $[t - \tau_d, t]$. This value can be calculated by Equation 11.

However, to avoid an unnecessary amount of evaluations of this function, the MOI is added as a state variable $\phi(t)$ to the model. The rate of change of this variable is given in Equation 12 and is equal to the derivative of Equation 11.

$$\phi(t) = \int_{t-\tau_d}^{t} \frac{\alpha \cdot P(t_i) \cdot B(t_i)}{B(t_i)} dt_i = \int_{t-\tau_d}^{t} \alpha \cdot P(t_i)\, dt_i$$

( 11 )

$$\frac{d\phi(t)}{dt} = \alpha \cdot P(t) - \alpha \cdot P(t - \tau_d)$$

( 12 )
